# A small regulatory RNA controls antibiotic tolerance in *Staphylococcus aureus* by modulating efflux pump expression

**DOI:** 10.1101/2024.08.05.606701

**Authors:** Kam Pou Ha, Philippe Bouloc

## Abstract

*Staphylococcus aureus* is an opportunistic pathogen that poses a considerable burden to healthcare settings worldwide, aided by its ability to thrive in different environmental growth conditions and survive exposure to antibiotics. Small regulatory RNAs (sRNAs) are crucial in enhancing bacterial fitness by modulating gene expression in response to changing environmental conditions. We investigated the role of sRNAs in *S. aureus* antibiotic resistance and tolerance. By assessing the fitness of a library of sRNA mutants, we identified that RsaA sRNA is required for bacterial growth when exposed to low concentrations of fluoroquinolone, a class of antibiotics targeting DNA replication. We also found that in the absence of RsaA, *S. aureus* is less susceptible to β-lactam antibiotics, which act on the cell wall. RsaA has been reported to prevent the expression of MgrA, a master regulatory protein controlling the expression of efflux pumps. Here, we show that RsaA affects the sensitivity of *S. aureus* to fluoroquinolone and β-lactam antibiotics through MgrA. RsaA has two forms, a short one that is commonly referred to in RsaA studies, and a long form about twice the length, of which not much is known. Interestingly, our phenotype was restored only when complemented with the long form of the gene. This work demonstrates the role of regulatory RNAs in the adaptation of *S. aureus* to antibiotic resistance and highlights their value as potential therapeutic targets for manipulating individual sRNA responses to promote the efficacy of existing antibiotics.

## Introduction

Antibiotic resistance refers to the ability of bacteria to grow in antibiotic concentrations that are considered inhibitory, while antibiotic tolerance is the capacity of bacteria to survive exposure to bactericidal antibiotics for extended periods without necessarily acquiring genetic changes (1). Resistance involves the acquisition of resistance genes from other bacteria or by genetic mutation (1–3), leading to mechanisms such as enzymatic inactivation of antibiotics, alteration of antibiotic targets, or efflux pumps that expel antibiotics from bacterial cells (2, 3). However, the molecular basis for tolerance is wider in range. Antibiotic tolerance can be mediated by the induction of stress responses, such as the SOS DNA damage response that detects and repairs damage caused by fluoroquinolone antibiotics (4, 5), or by the reversible depletion of peptidoglycan in the cell wall, which enables survival in the presence of β-lactam antibiotics (6). In an extreme subset of tolerance, also called persistence, bacterial cells may enter a dormant or near-dormant state to protect cellular processes against the action of antibiotics (7). Tolerance has also been demonstrated as a precursor to the development of antibiotic resistance (8–10).

Although antibiotic resistance mechanisms have been well studied, less is known about antibiotic tolerance. In addition, while global efforts have primarily focussed on methicillin-resistant *S. aureus* (MRSA), these strains represent only a subset of the overall *S. aureus* burden (11). Indeed, infections from both susceptible and resistant *S. aureus* strains lead to prolonged illness and increased mortality rates (11, 12). Therefore, understanding how *S. aureus* modulates its sensitivity to antibiotics in the absence of resistance could aid in the development of novel treatment options to prevent resistance from developing.

Small non-coding RNAs (sRNAs) are key players in the post-transcriptional regulation of target genes (13). They enable rapid environmental adaptation by base-pairing with their target mRNAs to modulate mRNA translation and stability (13). sRNAs act on almost all cellular functions and as a result, have emerged as important regulators of virulence (14), metabolic adaptation (15) and antibiotic resistance (16, 17). Although many sRNAs have been identified in *S. aureus,* their functions and targets are still mostly unknown (18, 19). The best-described sRNA in *S. aureus* is RNAIII, a large 514-nucleotide sRNA that is involved in virulence (20). Other described sRNAs include RsaE, which is associated with the TCA cycle (21, 22) and arginine catabolism (15), and IsrR, which plays a key role in the iron-sparing response (23, 24). Although sRNAs play important roles in regulating antibiotic-resistance processes such as cell envelope modifications and drug efflux pumps in Gram-negative species (16, 25–27), their impact on antibiotic resistance in Gram-positive species has received less attention (16, 17). In addition, the role of sRNAs in antibiotic tolerance is poorly documented.

To find sRNAs required for *S. aureus* to adapt to sublethal antibiotic concentrations, we used a fitness assay with sRNA mutants grown in the presence of the fluoroquinolone antibiotic norfloxacin. We identified a single sRNA, RsaA, the loss of which increased sensitivity of the bacterium to fluoroquinolone antibiotics. Further tests revealed that loss of RsaA also decreased the sensitivity of *S. aureus* to β-lactam antibiotics, and that both effects were mediated by RsaA via the master regulatory protein MgrA. Together, our results demonstrate that sRNAs contribute towards antibiotic tolerance in *S. aureus*.

## Results

### RsaA is required for growth in the antibiotic norfloxacin

To determine whether sRNAs are required for *S. aureus* to adapt to the presence of the fluoroquinolone antibiotic norfloxacin, we employed a well-established technique that enables the competitive fitness of a sRNA mutant library to be assessed (18, 23, 28, 29). This technique, referred to as the fitness competition assay, uses a library of DNA-tagged deletion mutants in the HG003 strain (Supplementary Table S1) and enables the subtle phenotypes of sRNA mutants to be screened more easily. Three mutants were independently constructed for each sRNA gene, which were then used to establish three independent libraries (Supplementary Table S4). Each library was restricted to 48 mutants of “*bona fide*” sRNAs, defined as those that are genetically independent with their own promoter and terminator, to prevent any interference from nearby coding sequences (18). sRNA gene sequences were replaced with specific DNA tags to enable each mutant to be identified and the proportion of each mutant within the total population to be calculated. By using indexed PCR primer pairs, up to 40 samples could be tested in one DNA-seq run (18). Mutants that disappeared or accumulated under a given stress condition indicated a potential role for the corresponding sRNAs under this growth condition.

The sRNA libraries were exposed to a sub-lethal concentration of norfloxacin and grown over three days with two serial dilutions. Samples were taken at different time points and growth stages, and the proportion of each mutant within the population was determined (Figure 1A). Results were normalised to the same inoculum grown in the same medium without antibiotic and sampled at the same growth phases. Among the 48 tested mutants, the strain containing Tag075 displayed the greatest fitness disadvantage, decreasing by 28-fold [log2(FC)=- 4.8] in the presence of norfloxacin after 72 hours, when compared to the control condition (Figure 1B). This strain, in which the *rsaA* gene has been replaced by Tag075 (*ΔrsaA*), progressively disappeared from the library under the selective pressure of norfloxacin under sub-lethal concentrations. Neither *rnaIII*, *rsaC*, *rsaD*, *rsaE*, *rsaG, isrR* nor *rsaOG* mutants (18), present in the libraries tested, were impacted by the presence of norfloxacin, indicating that their corresponding sRNAs are unlikely to contribute to adaptation to norfloxacin under the tested conditions.

**Figure 1.**
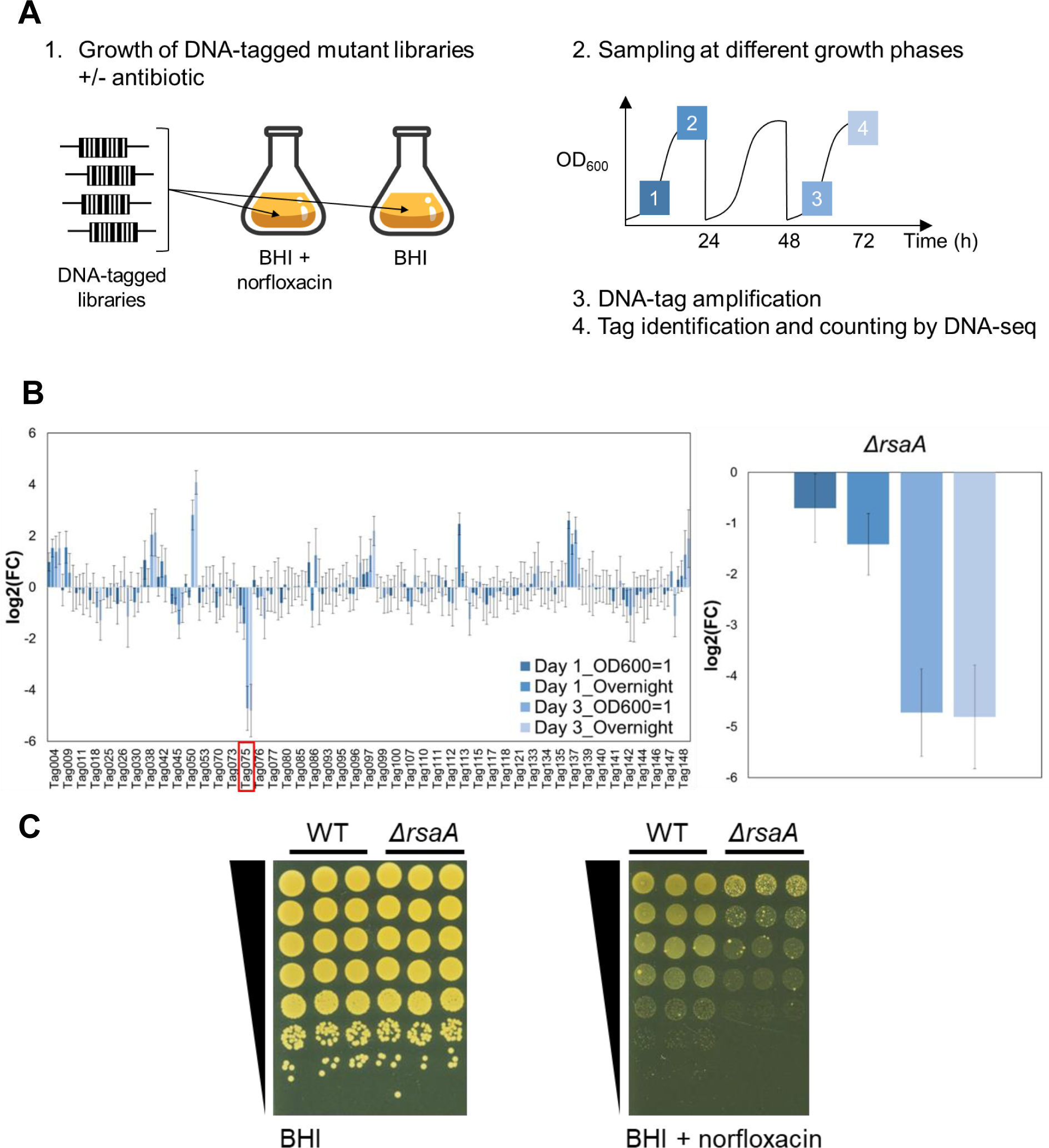
*S. aureus* RsaA is required for optimal growth in the presence of norfloxacin. **(A)** Experimental protocol scheme to identify mutants with altered fitness in media containing the antibiotic norfloxacin. **(B)** Evolution of mutant proportions in libraries (for composition, see Supplementary Table S4) grown in the presence of 0.25 µg/ml norfloxacin normalised to the same libraries grown in the absence of antibiotic. Error bars indicate the standard deviation from three independent libraries. Left: data with the complete 48 mutant libraries. Tag075 corresponding to the *rsaA* mutant is highlighted by a red box. For Tag-mutant association, see Supplementary Table S1. Right: selected data for *ΔrsaA* is shown. Colour code corresponds to the sampling time colour from Figure 1A. **(C)** Growth of WT and *ΔrsaA* strains serially diluted 10-fold and spotted onto BHI agar +/- 0.6 µg/ml norfloxacin (n = 3).

To assess whether the sensitivity of the *rsaA* mutant to norfloxacin also occurred in the absence of the 47 other sRNA strains, monocultures of *ΔrsaA* were grown on agar plates containing norfloxacin. Results from these spot tests revealed that *ΔrsaA* is 100-fold more sensitive to norfloxacin, when compared to its parental strain HG003, referred to here as the wild-type strain (WT) (Figure 1C). In contrast, no difference in growth was observed between either strain when grown in the absence of the antibiotic. Therefore, loss of RsaA leads to greater sensitivity of *S. aureus* to norfloxacin and this is independent of any effects on growth in a mixed culture.

Taken together with the fitness competition results, these findings suggested that the RsaA sRNA is required for optimal growth of *S. aureus* in the presence of norfloxacin.

### *S. aureus rsaA* norfloxacin-dependent phenotype is complemented only by the long form of the *rsaA* gene

To confirm that the increased sensitivity of the *rsaA* mutant to norfloxacin was solely due to the loss of RsaA, the *rsaA* mutant was complemented with a wild-type copy of the gene. Since there are two versions of the RsaA transcript, a short form of 144 base pairs in length and a long version of 282 base pairs (30, 31) (Figure 2A; Supplementary Figure S1), individual plasmids were constructed to reflect complementation with each form of the gene.

**Figure 2.**
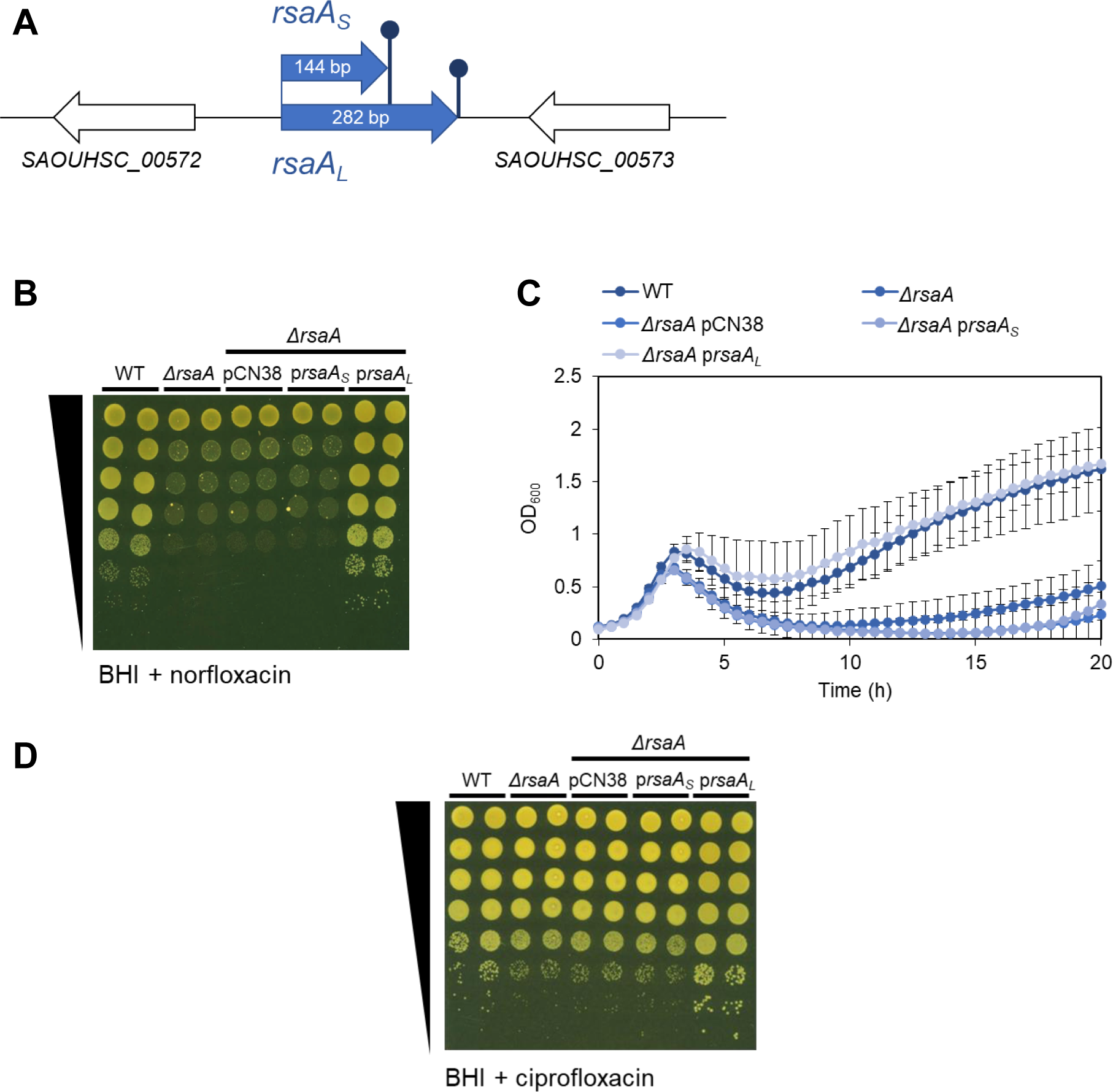
The long form of RsaA restores the wild-type phenotype, but not the short form. **(A)** Schematic diagram of long and short forms of RsaA in *S. aureus*. Blue and white arrows represent genes, dark blue sticks represent transcription terminators. Gene annotations refer to NCTC8325 nomenclature (CP00025.1) retrieved from Genbank (33). The predicted secondary structure of RsaA_L,_ along with the experimentally confirmed structure of RsaA_S_ (31), can be found in Fig. S1. **(B)** Growth of WT, *rsaA* mutant, empty vector (pCN38) and mutants complemented with RsaA short form (p*rsaA_S_*) or long form (p*rsaA_L_*), serially diluted 10-fold and spotted onto BHI agar + 0.6 µg/ml norfloxacin (n = 3). Spot tests in BHI broth alone are shown in Fig. S2. **(C)** Growth curves of WT and *rsaA* mutants grown in BHI broth containing 0.5 µg/ml norfloxacin (n = 3). OD_600_ measurements in BHI broth alone are shown in Fig. S2. Error bars represent the standard deviation of the mean. **(D)** Growth of WT, *rsaA* mutant, empty vector (pCN38) and mutants complemented with RsaA short form (p*rsaA_S_*) or long form (p*rsaA_L_*), serially diluted 10-fold and spotted onto BHI agar + 0.15 µg/ml ciprofloxacin (n = 3).

When the strains were spotted onto agar containing norfloxacin, the loss of RsaA reduced the growth of *S. aureus* by 100-fold when compared to the wild-type strain, but no improvement in growth occurred when the *rsaA* mutant was complemented with the short form of the gene (RsaA_S_) (Figure 2B), indicating that RsaA_S_ is not enough to restore the wild-type phenotype. This phenotype was also observed when the same strains were grown in liquid media containing norfloxacin (Figure 2C), with the difference that growth of *ΔrsaA* was 3.12-fold slower than the wild-type strain after 20 h rather than 100-fold, which reflects the higher sensitivity of spot tests when compared to growth curves. However, in both solid and liquid cultures, complementation of *ΔrsaA* with the long form of RsaA (RsaA_L_) restored the wild-type phenotype, demonstrating that the long form of the gene is essential for optimal growth of *S. aureus* in this antibiotic.

### Loss of RsaA sensitises *S. aureus* to ciprofloxacin

Norfloxacin belongs to the fluoroquinone class of antibiotics that cause DNA damage by blocking DNA replication (32), which leads to cell death. To determine whether the phenotype was specific to norfloxacin or could also be observed in another antibiotic of the same functional class, the spot test experiment was repeated with ciprofloxacin. As observed in Figure 2D, the *rsaA* mutant was more sensitive to ciprofloxacin than the WT strain, and this phenotype was restored when complemented with the long form of the gene. Therefore, RsaA_L_, provided here in multicopy form through a plasmid, leads to a phenotype that affects fluoroquinlones in general.

Combined, these data indicate that RsaA is involved in protecting *S. aureus* against the action of fluoroquinolone antibiotics, and that this function requires the long form of the gene to be present.

### Loss of RsaA reduces sensitivity of *S. aureus* to β-lactam antibiotics

Having demonstrated that RsaA is required for optimal growth of *S. aureus* in the presence of fluoroquinolone antibiotics, we tested whether a similar effect could occur for other antibiotic classes. For this, the β-bactam and glycopeptide antibiotic classes, represented by cefazolin and vancomycin respectively, were selected for testing due to their clinical relevance.

When the strains were grown in the presence of these two antibiotics, it was revealed that vancomycin had no effect on the growth of the *rsaA* mutant when compared to the wild-type strain (Figure S3). By contrast, in the presence of cefazolin, *S. aureus* growth was enhanced in the absence of RsaA by 10-fold in spot tests (Figure 3A) and by 1.2-fold in liquid culture after 20 h (Figure 3B), when compared to the wild-type strain. Furthermore, the wild-type phenotype was restored only when *ΔrsaA* was complemented by the long form of the gene.

**Figure 3.**
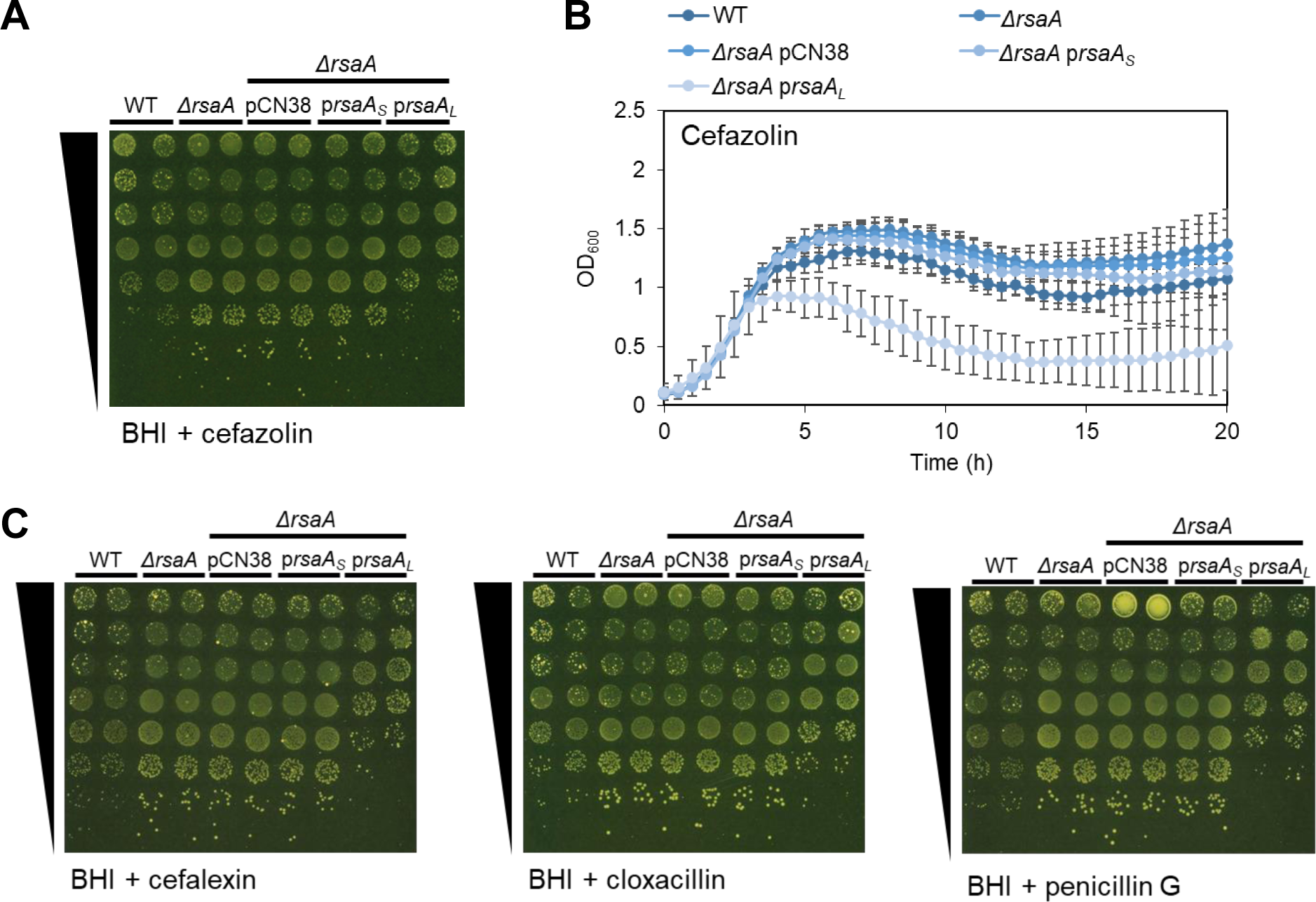
The *rsaA* mutant is less sensitive to β-lactam antibiotics when compared to the WT. **(A)** Growth of WT, *rsaA* mutant, empty vector (pCN38) and mutants complemented with RsaA short form (p*rsaA_S_*) or long form (p*rsaA_L_*), serially diluted 10-fold and spotted onto BHI agar + 0.3 µg/ml cefazolin (n = 3). Spot tests in BHI broth alone are shown in Fig. S2. **(B)** Growth curves of WT and *rsaA* mutants grown in BHI broth containing 0.125 µg/ml cefazolin (n = 3). OD_600_ measurements in BHI broth alone are shown in Fig. S2. Error bars represent the standard deviation of the mean. **(C)** Growth of WT, *rsaA* mutant, empty vector (pCN38) and mutants complemented with RsaA short form (p*rsaA_S_*) or long form (p*rsaA_L_*), serially diluted 10-fold and spotted onto BHI agar + 0.6 µg/ml cefalexin, 0.15 µg/ml cloxacillin or 0.03125 µg/ml penicillin G (n = 3). Spot tests in BHI broth alone are shown in Fig. S2.

As a follow up, we repeated the spot tests on a selection of other β-bactam antibiotics: cefalexin, cloxacillin and penicillin G. Results demonstrated that growth of the *rsaA* mutant is increased in the presence of β-bactams when compared to the wild-type strain (Figure 3C). Taken together, these data demonstrate that the loss of RsaA reduces the sensitivity of *S. aureus* to β-bactam antibiotics. This is opposite to the phenotype found for fluoroquinolone antibiotics, in which the loss of RsaA made *S. aureus* more sensitive to these antibiotics instead. Nevertheless, in both cases, the long form of RsaA is required for this process.

### RsaA acts on MgrA to fully mediate its effect on norfloxacin and partly for cefazolin

RsaA is known to repress translation of the master regulatory protein, MgrA (31). The latter regulates *S. aureus* virulence by promoting genes involved in capsule synthesis and inhibiting genes related to biofilm formation (31). Separately, MgrA has also been shown to act on several efflux pumps, including the fluoroquinolone efflux pumps NorA (34, 35), NorB (35–37) and NorC (38), and a β-bactam efflux pump, AbcA (37). Since MgrA is reported to inhibit these Nor efflux pumps and promote AbcA (34–38), this would align with our observed phenotypes in the *rsaA* mutant, in which MgrA is no longer being repressed. To test this hypothesis, we performed spot tests on single and double mutants of *rsaA* and *mgrA* in the presence of norfloxacin or cefazolin. As seen in Figure 4A, loss of RsaA and MgrA led to opposite effects on *S. aureus* growth when compared to the wild-type strain, supporting previous reports of RsaA as a repressor of MgrA. While *ΔrsaA* was more sensitive to norfloxacin than wild-type *S. aureus*, the *mgrA* mutant showed reduced sensitivity, and the reverse was true of these two strains concerning cefazolin. When the double mutant was tested, its growth in norfloxacin was restored to wild-type levels, indicating that the increased sensitivity of *ΔrsaA* was most likely caused by derepression of MgrA. In the presence of cefazolin, growth of *ΔrsaA ΔmgrA* was very slightly increased compared to *ΔmgrA* but does not reach *ΔrsaA* levels, again indicating that the phenotype is likely due to the derepression of MgrA. However, in this case, growth also did not reach wild-type levels, suggesting a minor role for another RsaA-mediated mechanism.

**Figure 4.**
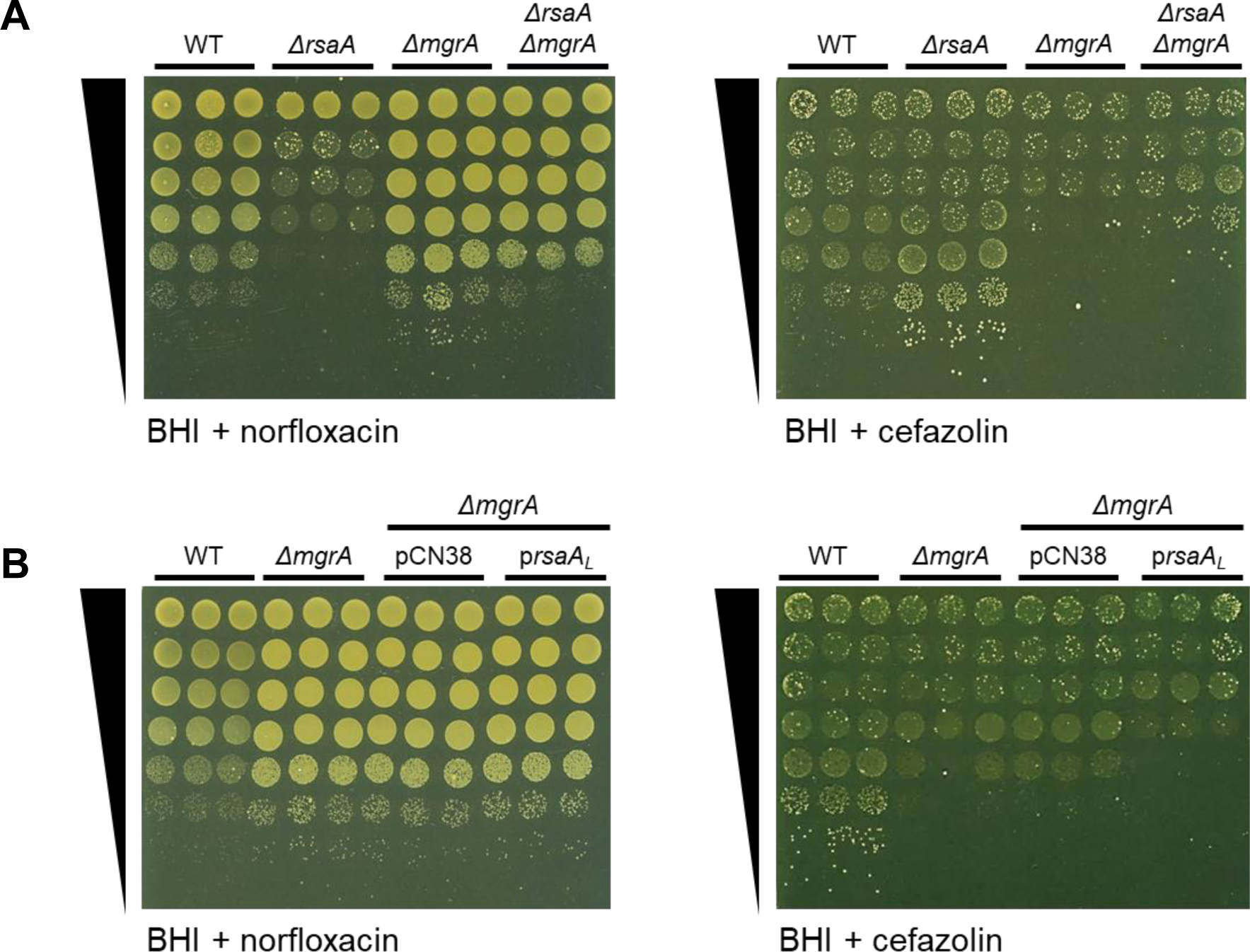
The increased sensitivity of the *rsaA* mutant to norfloxacin is mediated by MgrA. Growth of WT, *ΔrsaA*, *ΔmgrA* and *ΔrsaA ΔmgrA* strains, serially diluted 10-fold and spotted onto BHI agar containing **(A)** 0.6 µg/ml norfloxacin or **(B)** 0.3 µg/ml cefazolin (n = 3). **(B)** Growth of WT, *ΔmgrA* and *ΔmgrA* complemented with empty vector (pCN38) or long form (p*rsaA_L_*), serially diluted 10-fold and spotted onto BHI agar containing **(C)** 0.6 µg/ml norfloxacin or **(D)** 0.3 µg/ml cefazolin (n = 3). Spot tests in BHI alone are shown in Fig. S4.

To further investigate the link between RsaA, the derepression of MgrA and its subsequent effect on *S. aureus* sensitivity to norfloxacin and cefazolin, we generated an *mgrA* mutant that overproduced RsaA_L_ (*ΔmgrA* pRsaA_L_) and repeated the spot tests (Figure 4B). These results revealed that the overproduction of RsaA_L_ has no additional effect on the growth of *ΔmgrA* in the presence of norfloxacin, demonstrating that RsaA acts solely through MgrA to mediate its effect on norfloxacin. By contrast, the overproduction of RsaA_L_ led to a slightly increased sensitivity of *ΔmgrA* to cefazolin, indicating that while RsaA mostly acts through MgrA to mediate its effect on cefazolin, another mechanism may be involved.

### The full-length *rsaA* gene is required for expression of the short RsaA form

Having confirmed that the long form of RsaA was essential for mediating the sensitivity of *S. aureus* to growth in norfloxacin and cefazolin, we wanted to determine why this long form was required. To investigate this, we first searched for possible open-reading frames in the RsaA_L_ transcript that could translate to a small protein with a regulatory function. This identified one candidate that spanned most of the long-form region; however, mutation of the predicted ORF to block expression had no effect on antibiotic phenotype in spot tests (Supplementary Figure S5). Next, we examined whether additional interactions are present between *mgrA* mRNA and RsaA_L_, as compared to what is already known with *mgrA* mRNA and RsaA_S_ (31). However, RNA-RNA interaction prediction software did not find any new interactions present in RsaA_L_ that was not also detected with the short form only (Supplementary Figure S6), supporting that for RsaA to pair with *mgrA* mRNA, only its short form transcript is required.

Finally, we examined the levels of expression between RsaA_S_ and RsaA_L_. Results showed that there are far higher levels of the short form transcript in the wild-type strain when compared to the long form transcript (Figure 5A). This pattern is also reflected in the *rsaA* mutant when complemented with the long form of the gene (Figure 5A), corresponding with our previous observation that only RsaA_L_ restores the wild-type phenotype. However, when the *rsaA* mutant was complemented with RsaA_S_, minimal expression of RsaA_S_ was detected (Figure 5A), suggesting that while only the short form is required for sRNA function (Supplementary Figure S6), the short form cannot be expressed successfully on its own and expression of the long-form transcript is still needed. When we probed the same Northern blot membranes to detect only RsaA_L_ and the long-form region, it was revealed that for the complemented *rsaA* mutant that overexpressed RsaA_L_, we could also see a clear band corresponding to the long-form region only (Figure 5B), suggesting a cleavage of the long-form region from the RsaA_L_ transcript.

**Figure 5.**
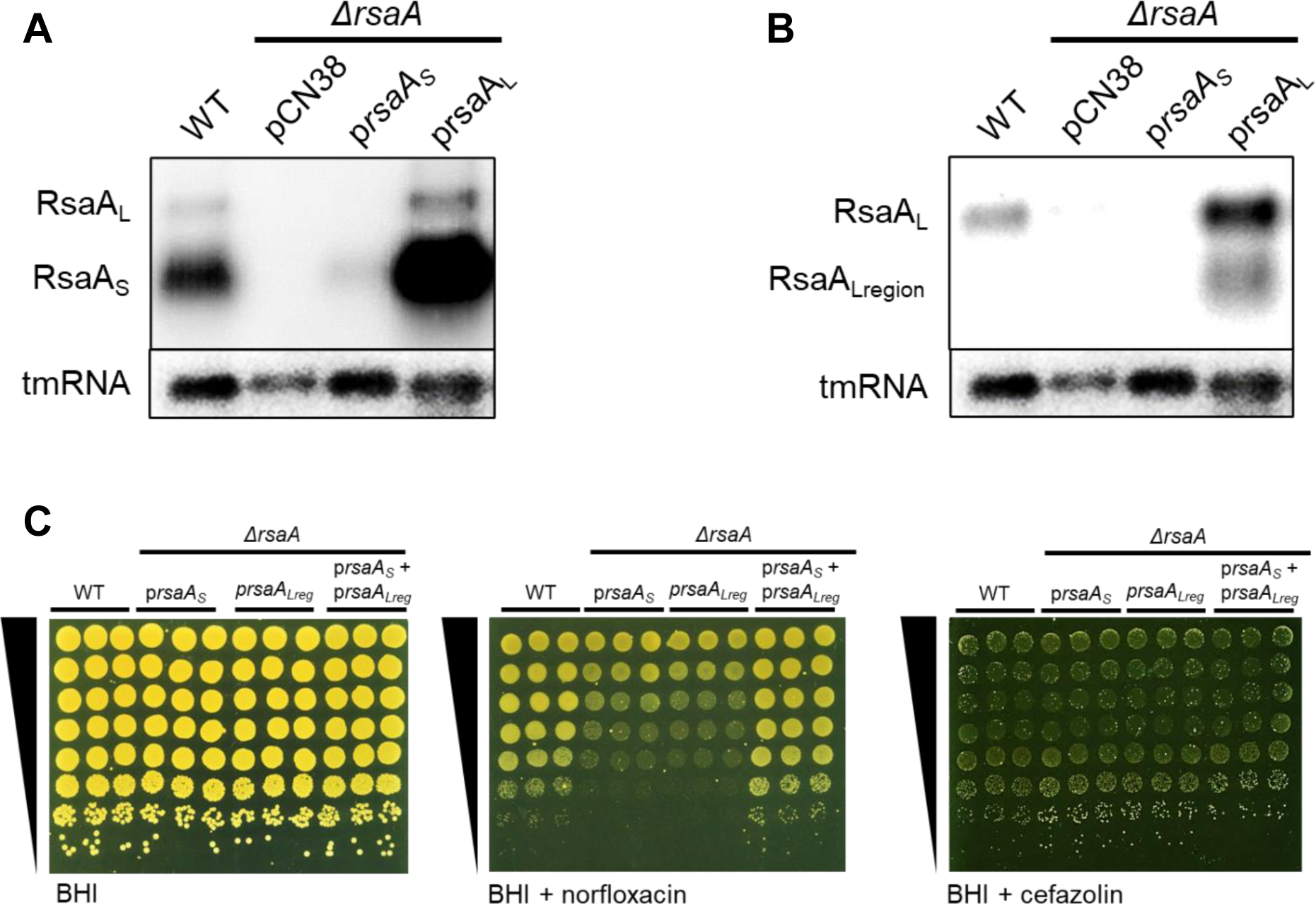
The long RsaA transcript needs to be expressed for the sRNA to function. **(A)** Northern blot of WT and *ΔrsaA* complemented with empty vector (pCN38), RsaA short form (p*rsaA_S_* or long form (p*rsaA_L_*), sampled at an OD_600_ of 6 and probed for RsaA using a 5’-end probe to detect both short and long forms, with tmRNA probed as a loading control (n = 4). **(B)** Northern blot using the same membrane as for Fig. 5A, probed for RsaA_L_ and RsaA_L-region_ only using a RsaA_L_ 3’-end probe, and tmRNA (loading control) (n = 4). **(C)** Growth of WT and *ΔrsaA* complemented with short form (p*rsaA_S_*), long-region only (p*rsaA_Lreg_*), or both short form and long region (p*rsaA_S_* + p*rsaA_Lreg_*), serially diluted 10-fold and spotted onto BHI agar +/- 0.6 µg/ml norfloxacin or 0.3 µg/ml cefazolin (n = 3).

To investigate the possibility of a cleavage of the long-form region further, the *rsaA* mutant was complemented with RsaA_S_ and only the long-form region (RsaA_L-region_) on separate plasmids. Spot tests revealed that although neither RsaA_S_ nor RsaA_L-region_ on their own can restore the wild-type phenotype, restoration of this phenotype occurs when both are present, even when they are not present on the same transcript (Figure 5C). This supports the hypothesis that the RsaA_L_ transcript is cleaved and that the two cleavage products are required for RsaA to function as a sRNA.

Combined, these data demonstrate that expression of RsaA_L_ is required for the short form to be generated, likely through a cleavage event in which both cleavage products are required for RsaA function.

## Discussion

Small regulatory RNAs play a crucial role in regulating key processes related to antibiotic resistance in Gram-negative species (16, 25). These processes include cell envelope modifications (26) and drug efflux pumps (27). However, their impact on antibiotic resistance in Gram-positive species has received less attention (16).

In *S. aureus,* the sRNA SprX, also known as RsaOR (21), influences vancomycin and teicoplanin glycopeptide resistance by repressing SpoVG (39). Notably, 6S RNA, which targets RNA polymerase, was shown to contribute to increased tolerance to low levels of rifampicin and related antibiotics in *S. aureus*, as well as in other distantly related bacteria (29). Antibiotics also influence sRNA gene expression, though connections to antibiotic resistance and/or tolerance are less clear. *S. aureus* sRNAs are differentially expressed after exposure to vancomycin, linezolid, ceftobiprole, and tigecycline (40), and exposure of *S. aureus* to sub-inhibitory concentrations of the β-lactam antibiotic oxacillin enhances bacterial virulence via the sRNA Ssr42 (41).

The data presented here connects the RsaA sRNA to antibiotic resistance mechanisms regarding to two antibiotic classes: fluoroquinolones (norfloxacin and ciprofloxacin) and β-lactams (cefazolin, cefalexin, cloxacillin and penicillin G). Although fluoroquinolones act on DNA replication and β-lactams on the cell wall, mechanisms of resistance to both classes can involve the action of efflux pumps. Our results show that RsaA acts through the master regulator MgrA to mediate the sensitivity of *S. aureus* to antibiotics, linking RsaA with the efflux pumps NorA, NorB, NorC and AbcA. To our knowledge, RsaA is the first example of sRNA linked to fluoroquinolone resistance mechanisms in *S. aureus*. Of note, in *E. coli*, the sRNA SdsR represses the TolC efflux pump and increases sensitivity to fluoroquinolones (27). SdsR activity is dependent on the RNA chaperon Hfq (27), which is likely not the case for RsaA (42).

We found that the effect of RsaA with fluoroquinolones was fully regulated via MgrA, but this regulation was only partial for β-lactams. Since sRNAs often control the expression of multiple mRNA targets to enable a coordinated regulation of cellular processes (13), it is likely that the RsaA-mediated effect on β-lactam sensitivity is mediated by other mRNA targets. A study investigating the RNA targetome of RsaA identified *ssaA_2* and *ssaA2_3* mRNAs as putative RsaA targets and found that these mRNAs form high-affinity complexes with RsaA *in vitro* (43). Since these two genes belong to the family of staphylococcal secretory antigen A (SsaA) proteins involved in peptidoglycan degradation (44), regulation of these substrates by RsaA may affect changes in sensitivity to β-lactam antibiotics, which function by disrupting peptidoglycan cross- linking during cell wall synthesis (45).

Loss of RsaA was complemented only by the long form of the gene, which demonstrates that the short form of the gene is not sufficient for regulating sensitivity to our tested antibiotics. RsaA_L_ was first detected by Northern blot and believed to originate from read-through at the RsaA_S_ transcriptional terminator (30). Both RsaA_L_ and RsaA_S_ share the same 5’ end, as determined by RACE experiments (22). Our study reveals the importance of the long-form *rsaA_L_* gene being present for the sRNA to function. As no additional interaction sites with *mgrA* mRNA were predicted in RsaA_L_ when compared to RsaA_S_, this correlates with studies on RsaA that have focussed only on its short form (30, 31, 43), which is believed to be the functional region. It must be noted that even if RsaA_L_ was not mentioned in these studies, the *rsaA* mutant was complemented using a long stretch of the *S. aureus* genome comprising of both *rsaA_L_* and *rsaA_S_* genes and the surrounding genomic region (30, 31, 43).

Our data demonstrate that the long form of the *rsaA* gene is needed for regulating staphylococcal sensitivity to norfloxacin and cefazolin, and that for RsaA to function, the long form transcript requires processing, most likely through cleavage into two halves. This finding aligns with measurements of the half-lives of RsaA short and long form transcripts (30), where RsaA_S_ was found to be > 2-fold more stable than RsaA_L_ after treatment with rifampicin to inhibit RNA polymerase (30). Based on experimental and homology data, there are believed to be 15 ribonucleases (RNases) in *S. aureus* that are involved in RNA processing and/or degradation (46). Of these, the main staphylococcal RNases are the exonucleases RNase J1/J2 and PNPase, and the endonucleases RNase III and RNase Y, the latter two of which have been reported to affect the stability and processing of RsaA_L_ (30, 47).

In *S. aureus,* the only other sRNA that is believed to be processed from a longer transcript is RsaE, which enables it to interact more efficiently with a second target, thereby expanding the range of targets recognised by this sRNA (48). In the case of RsaA, since we were able to restore the wild-type phenotype in a *rsaA* mutant by expressing both RsaA_S_ and the long region of RsaA_L_ from different plasmids, the RsaA_L_ long region is most likely interacting with the short form of RsaA to enable its function, possibly by stabilising the sRNA and/or by promoting interactions with *mgrA* mRNA.

Small regulatory RNAs control many cellular processes and enable rapid adaptation to stress. Here, we show that staphylococcal RsaA acts via the master regulator MgrA to mediate sensitivity to two different antibiotic classes, fluoroquinolones and β-bactams. In addition, we revealed that this regulation requires the full-length version of the *rsaA* gene, the transcript of which is likely processed by the cell to produce two halves that interact to produce functional RsaA. Therefore, exposure to sub-inhibitory fluoroquinolone or β-lactam concentrations can alter the expression of their corresponding efflux pumps, leading to changes in antibiotic tolerance.

## Methods

### Bacterial strains, plasmids and growth conditions

The bacterial strains used in this study are listed in Supplementary Table S1. Experiments were performed with HG003, a *Staphylococcus aureus* model strain widely used for regulation studies (49). Plasmids were engineered by Gibson assembly (50) in *E. coli* IM08B (51) as described (Supplementary Table S2), using the indicated appropriate primers (Supplementary Table S3) for PCR amplifications. Plasmids were verified by DNA sequencing and transformed into HG003 or its derivatives, as appropriate. HG003 *mgrA* and *rsaA mgrA* double mutants were constructed by transferring *mgrA*::tetM (52) into HG003 and HG003 *ΔrsaA*::tag075 using phage-mediated transduction.

*S. aureus* was routinely cultured in Brain Heart Infusion (BHI) broth at 37 °C, with shaking (180 rpm). *Escherichia coli* was grown in Lysogeny Broth (LB) at 37 °C with shaking (180 rpm). Media were supplemented with antibiotics as required: ampicillin 100 μg/ml for *E. coli*; chloramphenicol 5 μg/ml, tetracycline 1 μg/ml, and kanamycin 90 μg/ml for *S. aureus*.

### Fitness competition assay

Mutants with altered fitness in the presence of norfloxacin were identified and analysed using a reported strategy (28) with three independent libraries containing 48 mutants (Supplementary Table S4). The libraries were grown at 37°C in BHI or BHI + 0.25 µg/ml norfloxacin for 3 days. Overnight cultures were diluted 1000 times into fresh medium for each day. Samples were withdrawn at an OD_600_ of 1 and after overnight growth as indicated (Figure 1A).

### Spot tests

To assess mutant phenotypes, bacterial growth in the presence of antibiotics was visualised by spot tests, which enable subtle growth phenotypes to be detected, such as with sRNA mutants. 5 µL of ten-fold serial dilutions of overnight cultures (from neat to 10^-7^ dilution) were spotted on agar plates containing either BHI or BHI + antibiotic at the indicated concentrations. Agar plates were incubated for 24 h at 37 °C.

### Measurement of bacterial growth in liquid culture

To measure growth of *S. aureus* in liquid culture, bacteria were first grown to stationary phase in BHI at 37 °C (180 rpm), then inoculated 1/50 into a flat-bottomed 96-well plate (200 µl total volume) and placed into a CLARIOstar Omega plate reader (BMG Labtech). Antibiotics were added to the media as needed, at the indicated concentrations. Bacteria were grown for 20 h at 37 °C (300 rpm), and absorbance at 600 nm was measured every 30 min.

### Northern blots

Total RNA preparations and Northern blots were performed as previously described (53). 20 µg of total RNA samples were separated either by agarose (1.3%) gel electrophoresis. Membranes were probed with primers (Table S3) ^32^P-labelled using Terminal Deoxynucleotidyl Transferase (ThermoScientific) and scanned using an Amersham Typhoon imager.

## Acknowledgements

USA300 LAC *mgrA*::tetM strain used to transduce into HG003 to generate our *mgrA* mutants was a gift from Alexander Horswill. The research was supported by the “Agence National pour la Recherche” [ANR-19-CE12-0006-01 (sRNA-RRARE)].

